## Supplementary Information for "Acetylation-dependent coupling between G6PD activity and apoptotic signaling"

### Supplementary Figures

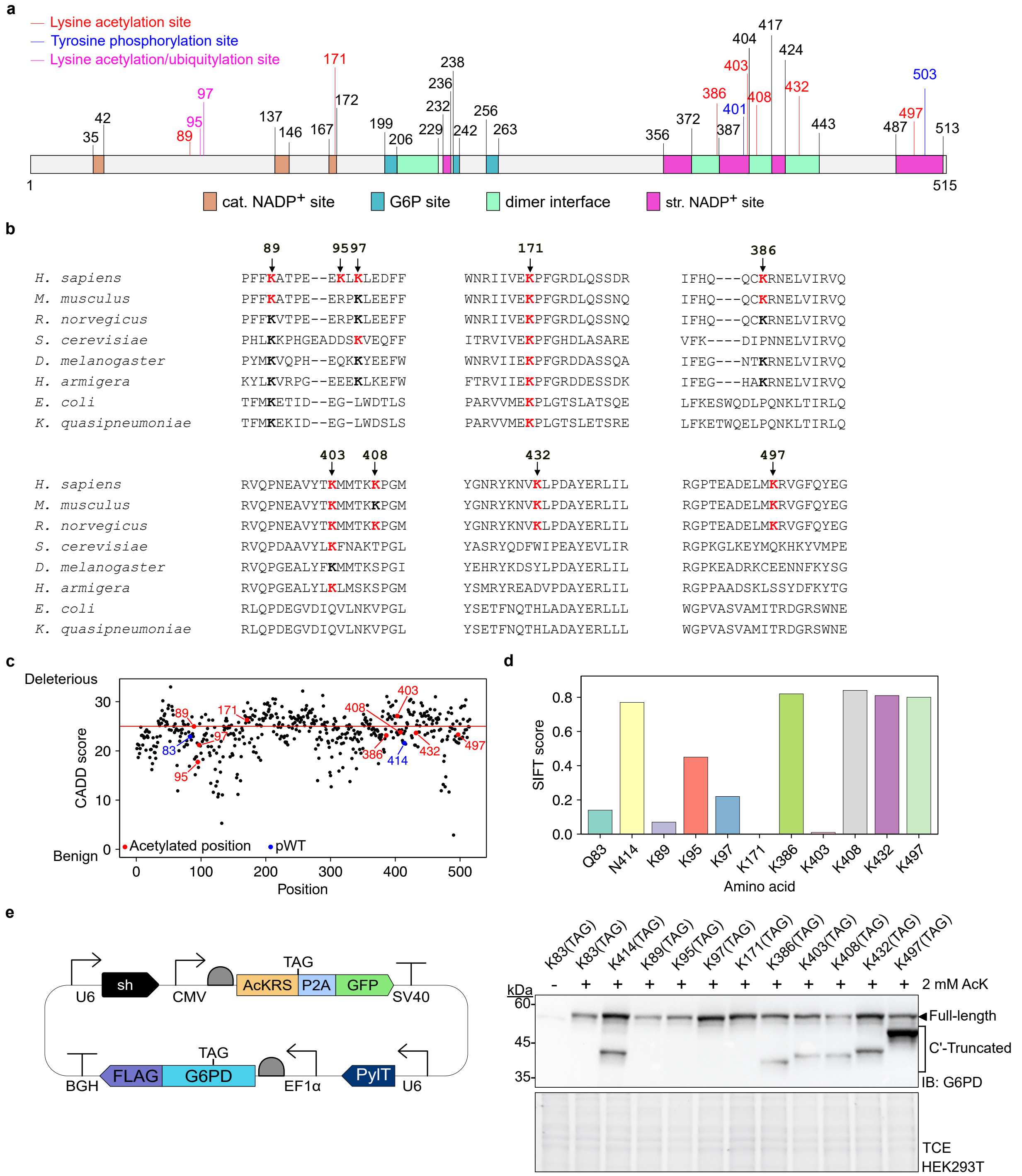


**Supplementary Fig. 1** | **a** Schematic representation of human G6PD primary sequence, its catalytic (cat.) NADP$^+$, structural (str.) NADP$^+$, and G6P binding sites, as well as its dimer interface. Posttranslational modification sites studied in this work are marked in red (lysine acetylation), blue (tyrosine phosphorylation), or magenta (lysine acetylation/ubiquitylation). **b** Multiple sequence alignment of G6PD from indicated organisms. Evolutionarily conserved lysine residues are marked in bold letters. Lysine residues, found to be acetylated by mass spectrometry-based proteomic studies, are highlighted in red. The numbering of acetylation positions selected for this study corresponds to human G6PD. **c** CADD score, which assigns a score between 35 (damaging) and 0 (benign), measuring variant deleteriousness that can effectively prioritize causal variants in genetic analyses[1]. The nine putative acetylation sites selected for this study are marked in red and pWT variants are marked in blue. **d** SIFT score, which uses evolutionary conservation, assigns a score between 0 (damaging) and 1 (benign)[2]. The analysis suggests that a lysine-to-alanine mutation at positions 89, 97, 171, and 403 is more harmful.

**Supplementary Fig. 1 Continued: e** Overexpression of acetylated G6PD in cultured mammalian cells. Left: Schematic description of the expression vector used for the simultaneous exogenous expression of acetylated G6PD and knockdown of endogenous G6PD. The vector includes the genes of an orthogonal acetylated lysine tRNA synthetase/pyrrolysine tRNA (AcKRS/$tRNA^{Pyl}_{CUA}$) pair for site-specific incorporation of acetylated lysine (AcK) in response to an in-frame UAG stop codon, shRNA against endogenous G6PD, and a C-terminally Flag-tagged (and sh-resistant) G6PD. The gene encoding the expression of AcKRS was cloned with a 3'-TAG stop codon followed by a self-splicing P2A sequence and green fluorescent protein (GFP), for UAG-suppression-dependent expression of GFP[3]. Eleven versions of this expression vector were initially cloned, in which an in-frame amber stop codon (TAG) mutation was introduced at a different position (83, 89, 95, 97, 171, 386, 403, 408, 414, 432, and 497). As controls, we also included a vector with the wild type (WT) G6PD gene (i.e., no TAG mutation) and a vector without the G6PD gene (termed sh). Right: Western blot shows acetyl lysine-dependent exogenous expression of acetylated G6PD variants in HEK293T cells. All eleven acetylated variants of full-length G6PD were expressed in the presence of 2 mM AcK, demonstrating that expression is AcK-dependent. Equal amounts of total protein were analyzed in each lane [as evident by 2,2,2-trichloroethanol (TCE) fluorescence]. Expression of C'-truncated G6PD was observed for K386TAG G6PD ($G6PD^{1\text{-}385}$), K403TAG G6PD ($G6PD^{1\text{-}402}$), K408TAG G6PD ($G6PD^{1\text{-}407}$), N414TAG G6PD ($G6PD^{1\text{-}413}$), K432TAG G6PD ($G6PD^{1\text{-}431}$), and K497TAG G6PD ($G6PD^{1-496}$). Source data are provided as a Source Data file.

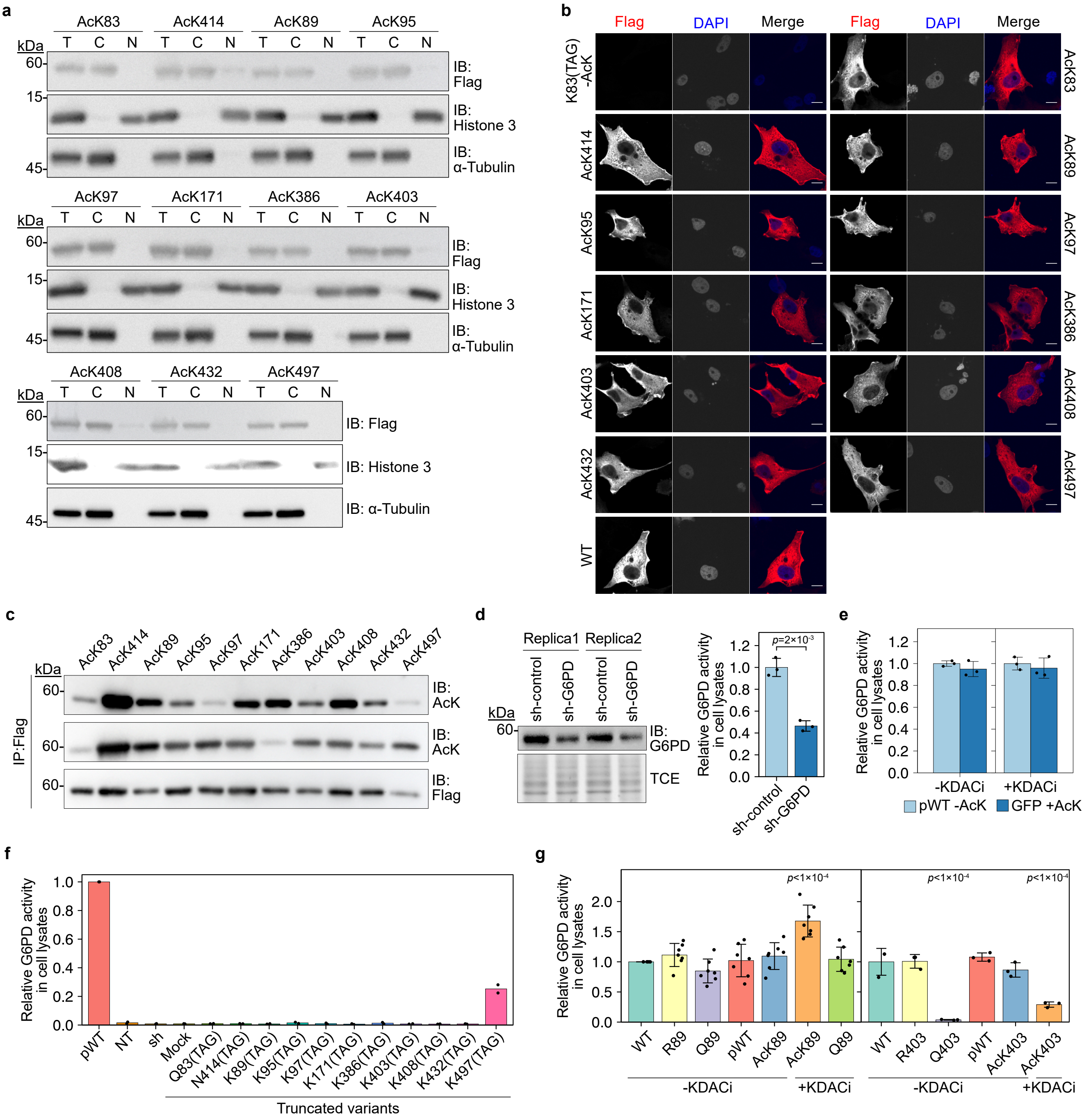


**Supplementary Fig. 2** | **a** Subcellular fractionation confirmed the cytoplasmic localization of acetylated G6PD and pseudo wild type (pWT) G6PD and in HEK293T cells. HEK293T cells transiently expressing pWT or indicated acetylated G6PD variants were treated with deacetylase inhibitors (KDACi). Total lysates (T), cytoplasmic (C), and nuclear (N) fractions were examined by immunoblot analysis using an anti-Flag antibody. Histone 3 and α-tubulin served as loading controls for the nuclear and cytoplasmic fractions, respectively.

**Supplementary Fig. 2 Continued:** **b** Representative confocal fluorescence microscopy images of COS7 cells expressing the indicated C-terminally Flag-tagged G6PD variants in the presence of KDACi. The ectopic protein was detected by immunofluorescence staining with Flag primary antibody. No apparent difference in cellular distribution was found between acetylated G6PD variants and WT G6PD. The scale bar represents 10 µm. **c** Western blots show differential recognition of site-specifically acetylated G6PD by commercially available anti-AcK antibodies. Indicated acetylated G6PD variants, expressed in HEK293T cells cultured with KDACi, were immunoprecipitated and detected by Western blotting using primary anti-AcK antibody from Cell Signaling (top) or Abcam (middle). **d** Knockdown of endogenous G6PD by encoded shRNA. Cells were transfected with plasmids encoding scrambled shRNA (sh-control) or shRNA against G6PD (sh-G6PD), and equal amounts of total cell lysate were used to evaluate G6PD expression levels (left), or to measure enzymatic activity under $V_{\text{max}}$ conditions (right). Data were analyzed using two-sided T-test and are presented as mean values ± SD; $n$=3 biologically independent samples. **e** Misincorporation, amber suppression, and KDACi have no effect on background G6PD activity measured in cell lysate. HEK293T cells expressing pWT G6PD were cultured without AcK to simulate the maximal level of background G6PD activity as a result of full-length G6PD expression due to misincorporation (UAG readthrough). In addition, HEK293T cells expressing GFP (as well as the amber suppression machinery) were cultured in the presence of AcK to simulate the effect of amber suppression on the catalytic activity of knocked-down endogenous G6PD. Enzymatic activity of G6PD was measured in cleared cell lysates and presented relative to the readthrough background activity. To verify that KDACi do not interfere with catalytic activity measurements performed in cell lysates or affect background G6PD activity, cells were cultured with (+) or without (-) KDACi. Data show that the possible expression of full-length G6PD due to misincorporation has negligible contribution to the background catalytic activity of knocked-down endogenous G6PD in cells with active UAG suppression machinery. This observation is independent of the presence of KDACi in culture media. Data are presented as mean values ± SD; $n$=3 biologically independent samples. **f** Enzymatic activity of C'-truncated G6PD. Truncated versions of G6PD were expressed in HEK293T cells cultured without AcK, and G6PD enzymatic activity was measured in cell lysates under $V_{\text{max}}$ conditions. Non-transfected cells (NT), G6PD knocked-down cells (sh), and cells expressing GFP (mock) were also included. Enzymatic activity of G6PD is presented relative to the activity measured in lysates of cells cultured with AcK and expressing full-length pWT G6PD. A measurable catalytic activity of C'-truncated G6PD was found only for G6PD$^{1-496}$ (K497TAG mutant of G6PD), which probably accumulates in cells together with AcK497 G6PD. Data are presented as mean values; $n$=2 biologically independent samples. **g** Enzymatic activity of K89-acetylated G6PD (left) or K403-acetylated G6PD (right) and mutants mimicking the acetylated and non-acetylated states. Wild type, pWT, AcK89, and AcK403 G6PD, as well as Lys-to-Arg or Lys-to-Gln mutants serving as mimics for the non-acetylated or acetylated states, respectively, were expressed in HEK293T cells cultured in the presence or absence of KDACi. Bars represent $V_{\text{max}}$ measured in cell lysates, normalized to $V_{\text{max}}$ of WT G6PD. The activity of G6PD K89R and K89Q mutants was similar to the activity of WT and pWT G6PD, while the activity of AcK89 G6PD (expressed in the presence of KDACi) was ∼1.7 fold higher. The K403R mutation had no effect on the catalytic activity of G6PD, but K403Q G6PD was inactive, and the activity of AcK403 G6PD was less than 30% of WT G6PD. These data show that lysine to glutamine mutation can mimic the effect of K403 acetylation, but not K89 acetylation, on G6PD enzymatic activity. Data were analyzed using one-way ANOVA followed by Tukey's post hoc test and are presented as mean values ± SD; $n$= 3 or 7 biologically independent samples. Source data are provided as a Source Data file.

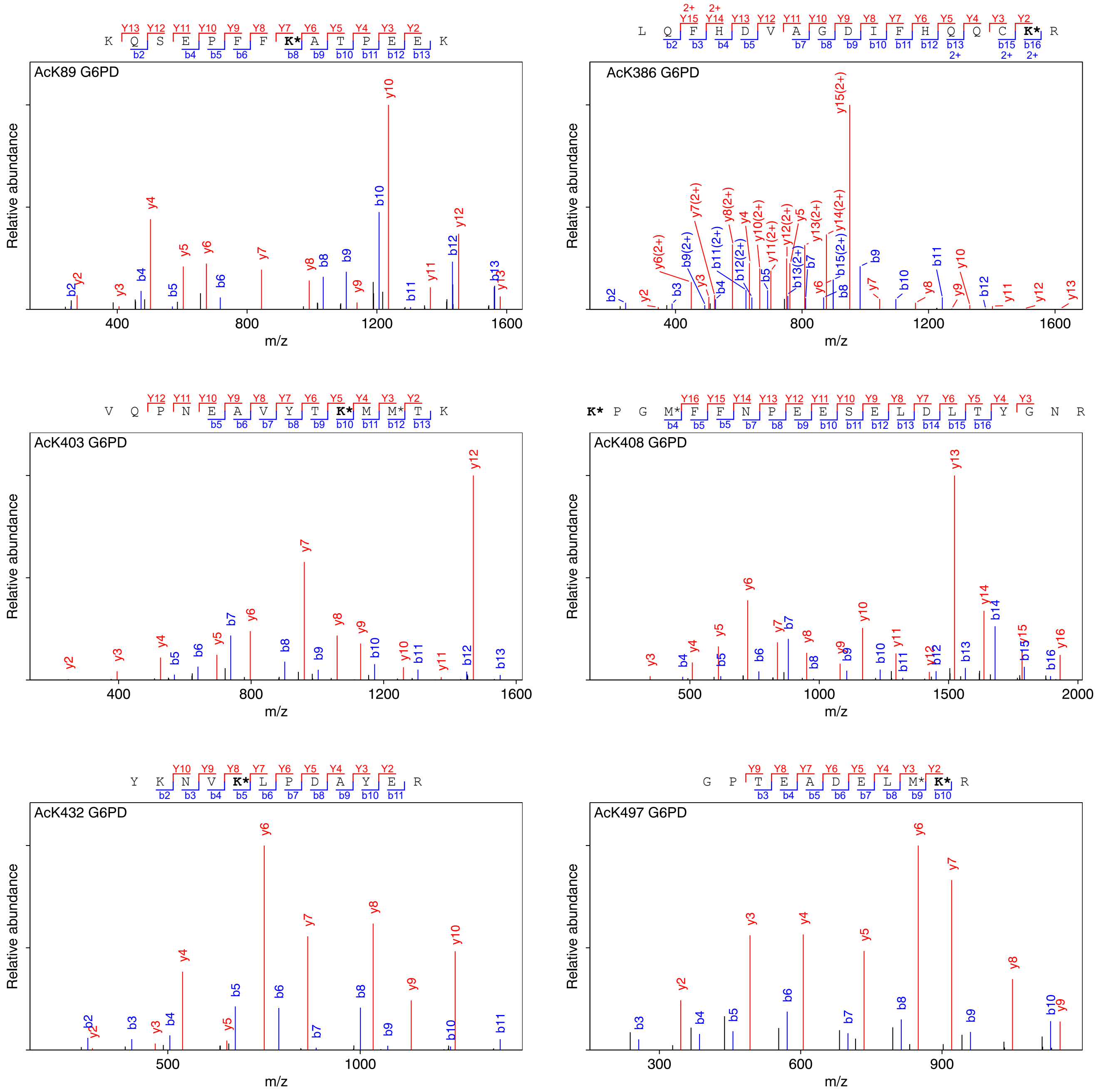


**Supplementary Fig. 3** | LC-MS/MS analyses of bacterially expressed and purified acetylated G6PD. The observed masses and sequences of presented tryptic peptides show the site-specific incorporation of AcK (indicated by **K***) into the designated positions. The modified peptide sequence and the precursor ion's measured mass are presented. Source data are provided as a Source Data file.

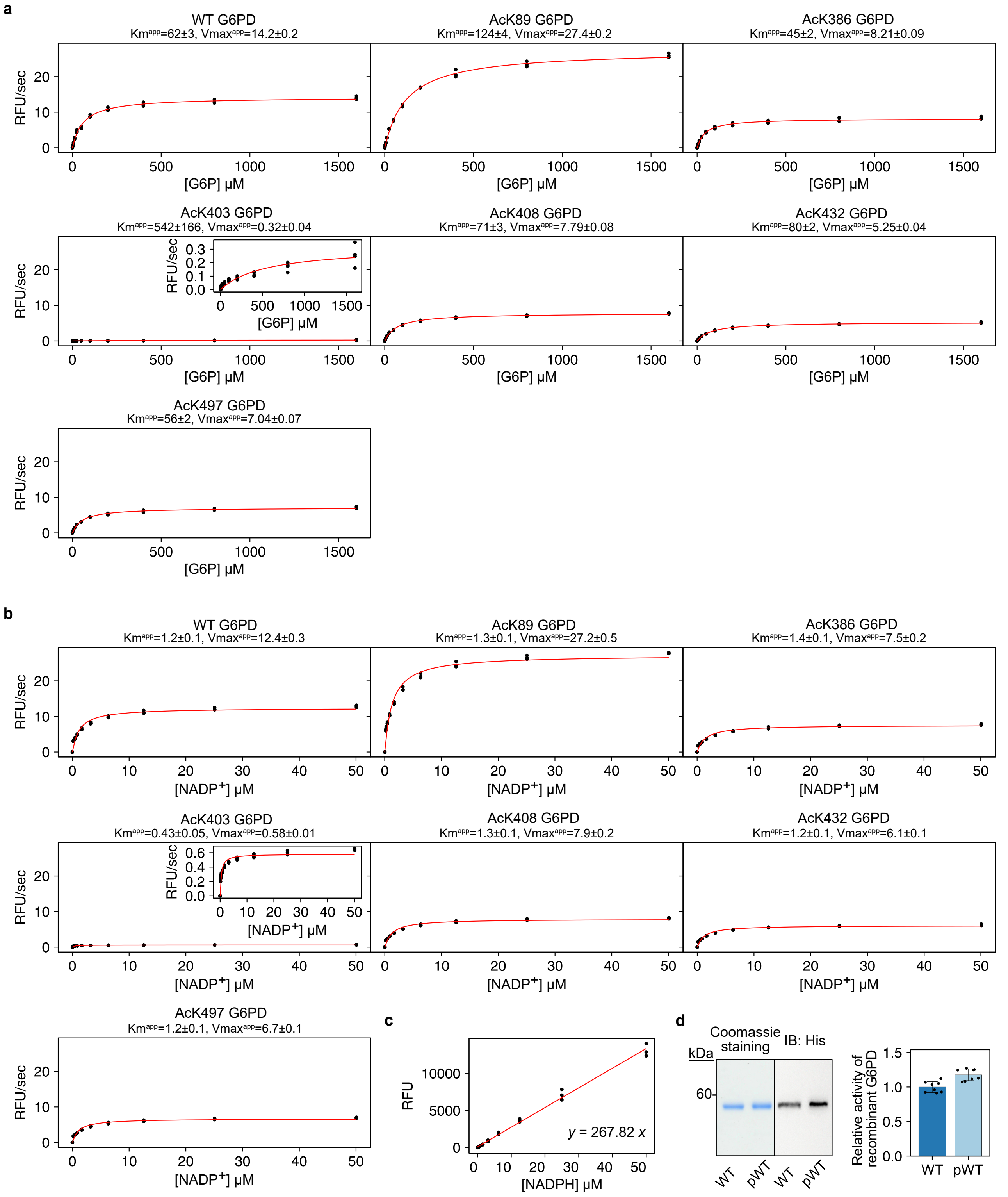


**Supplementary Fig. 4** | Steady-state kinetic analyses of G6PD enzymatic activity as a function of G6P concentration (**a**) or NADP$^+$ concentration (**b**). Apparent kinetic parameters ($V_{max}^{app}$ and $K_M^{app}$) were calculated by fitting the data to the Michaelis-Menten equation (red line). Values are reported ± fitting error, $n$=4 independent experiments. **c** Resorufin fluorescence as a function of initial NADPH concentration. Increasing concentrations of NADPH were incubated with diaphorase and an excess of resazurin. During the reaction, NADPH was consumed with the conversion of an equimolar amount of non-fluorescent resazurin to fluorescent resorufin. Endpoint measurements were fitted to a linear model, and the slope (267.82) was used to convert relative fluorescence units (RFU) presented in **a** and **b**, to $V_{max}$ values presented in Fig. 2b and Supplementary Table 2. $n$=3 independent experiments. **d** Equal amounts of bacterially expressed and purified WT and pWT G6PD were separated by SDS-PAGE and detected by Coomassie staining (left) or by immunoblotting using antibodies against the C-terminal 6×His tag (right). The relative catalytic activity of recombinant WT and pWT G6PD was measured under $V_{max}$ conditions. Data are the mean ± SD; $n$=8 independent experiments. Source data are provided as a Source Data file.

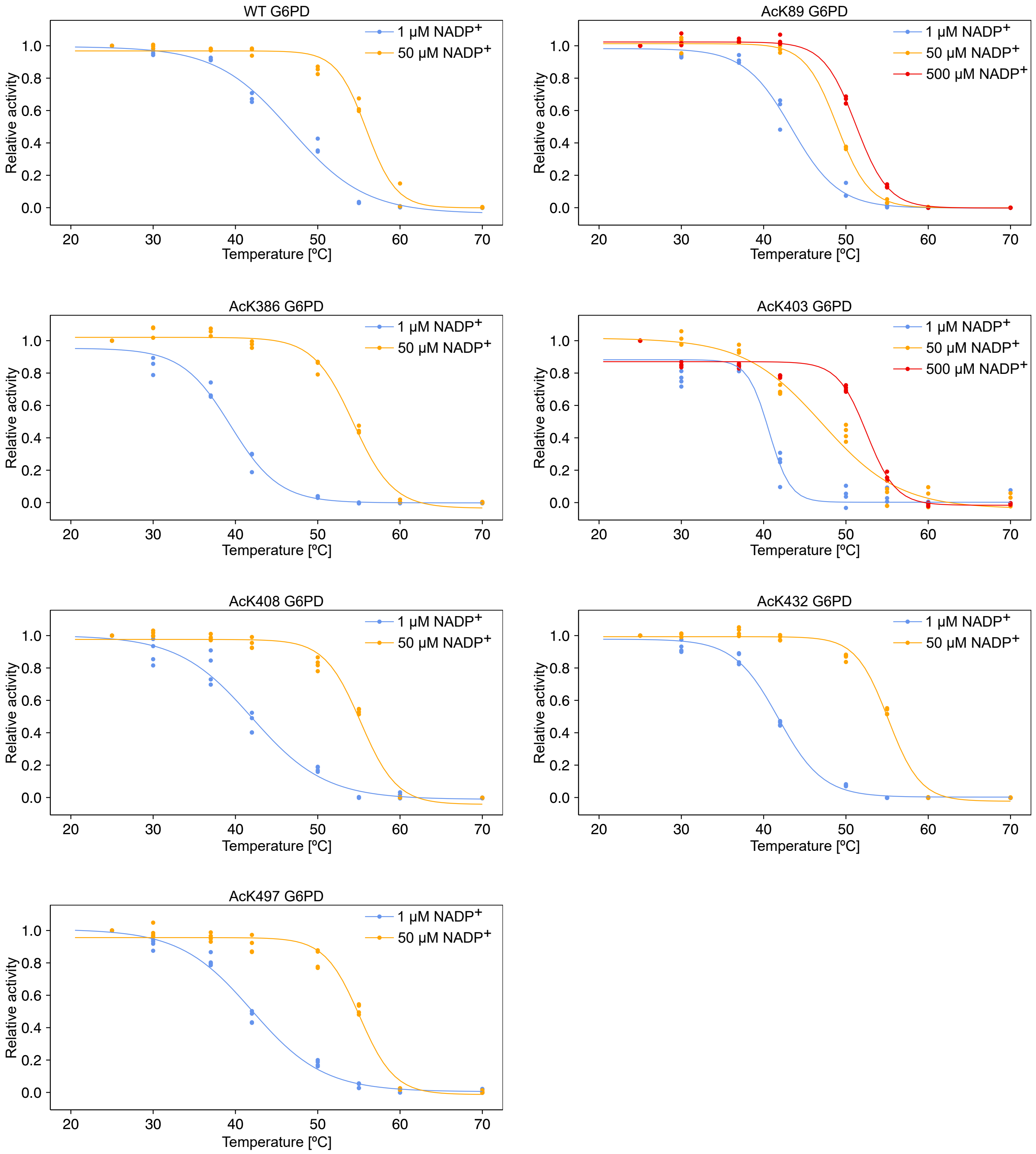


**Supplementary Fig. 5** | Thermal inactivation of WT and acetylated G6PD. Purified G6PD variants were incubated at increasing temperatures in the presence of indicated $NADP^+$ concentration, and G6PD enzymatic activity was then measured under $V_{max}$ conditions. Residual activity, relative to the activity measured following incubation at 25°C, was plotted as a function of incubation temperature and $NADP^+$ concentration. Lines represent fitting of data to a four-parameters logistic function from which $T_{1/2}$ values were extracted (summarized in Fig. 2d). $n$=3 or 4 independent experiments. Source data are provided as a Source Data file.

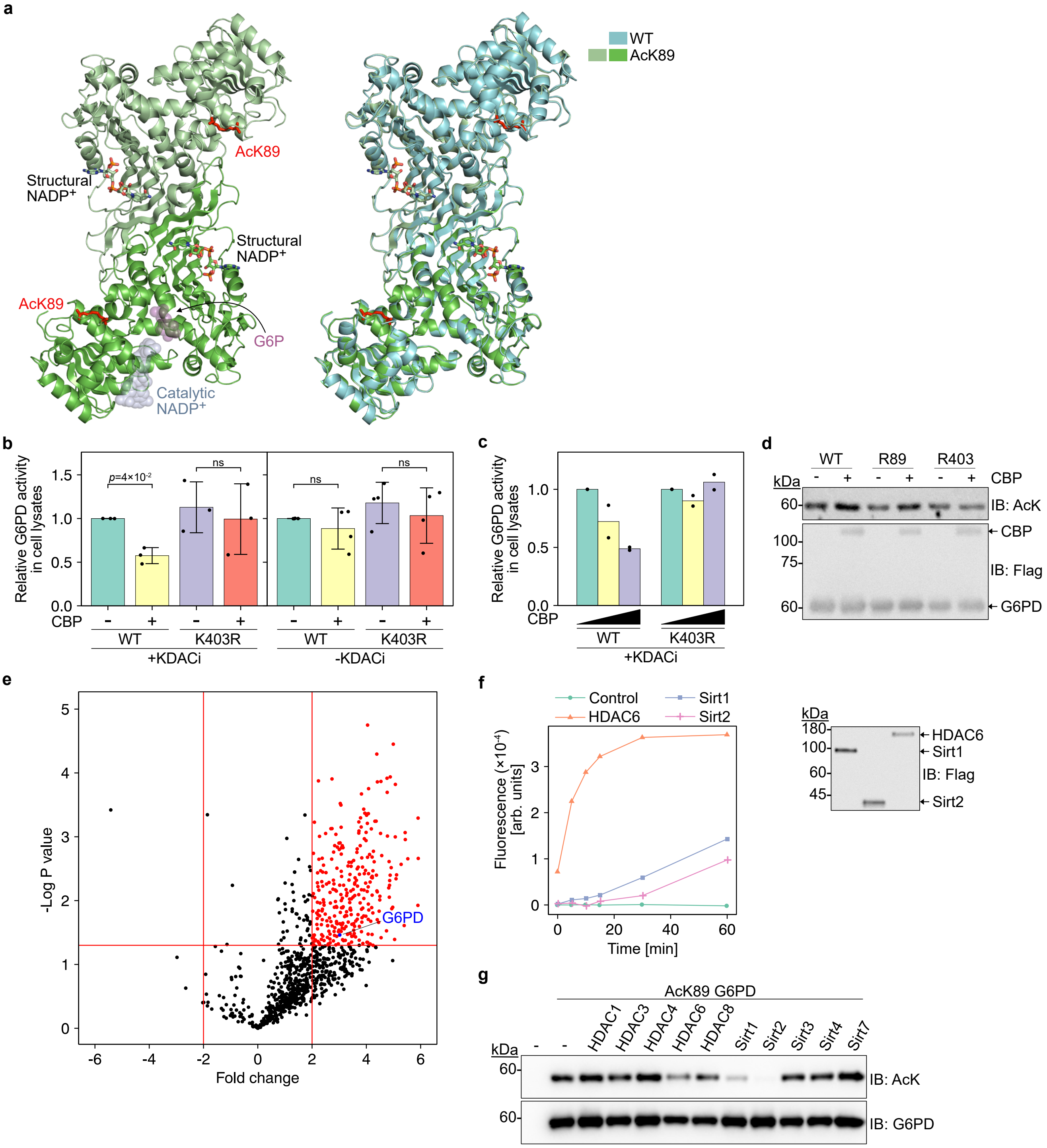


**Supplementary Fig. 6** | **a** Dimeric structure of AcK89 G6PD. Left: The dimeric structure of G6PD was constructed by symmetry operations, and the two monomers are presented in two shades of green. Acetylated K89 and structural NADP$^+$ are presented in sticks model. The expected positions of G6P (substrate) and catalytic NADP$^+$ are displayed as transparent spheres. Right: Superposition of dimeric AcK89 G6PD (green shades) and WT G6PD (cyan, PDB ID: 6E08), showing no significant change in G6PD dimeric structure following acetylation of position K89. **b** Lysine 403 is acetylated by CREB binding protein (CBP). HEK293T cells were seeded in a 24-well plate format and transiently transfected with 250 ng of the indicated G6PD expression plasmid together with 250 ng of CBP expression plasmid (+) or empty vector (-). Forty-two hours post-transfection, cells were treated with either KDACi or corresponding solvents for 6 h. Bars represent G6PD $V_{max}$ measured in cell lysates and normalized to G6PD protein level. Data were analyzed using one-way ANOVA followed by Tukey's post hoc test and are presented as mean values ± SD; $n$=3 (+KDACi), or 4 (-KDACi) biologically independent samples. Co-expression with CBP deactivated WT G6PD only in the presence of KDACi, suggesting deactivation by reversible acetylation. The inhibitory effect was not observed in the K403R mutant, indicating that position K403 of G6PD is a substrate of CBP.

**Supplementary Fig. 6 Continued: c** HEK293T cells were co-transfected with 250 ng of indicated G6PD expression plasmids, and increasing amounts of CBP-expressing plasmid (0, 250, and 500 ng). Forty-two hours post-transfection, cells were further treated with KDACi for 6 h. Bars represent the normalized G6PD $V_{\mathrm{max}}$ measured in cell lysates, relative to $V_{\mathrm{max}}$ measured in the absence of CBP ($n$=2 biologically independent samples). The decrease in $V_{\mathrm{max}}$ of WT G6PD as a function of CBP level was not observed in K403R G6PD, indicating that CBP can deactivate G6PD by acetylation of K403. **d** Western blot shows acetylation level of immunoprecipitated WT and mutants of G6PD. HEK293T cells were co-transfected with 250 ng of a plasmid expressing indicated G6PD variant and 500 ng of a plasmid expressing CBP or an empty vector, and cultured in the presence of KDACi. Acetylation level was detected by Western blotting. An increase in acetylation signal following co-expression with CBP was found in WT and K89R G6PD, but not K403R G6PD, suggesting that K403 G6PD is a substrate of CBP. **e** Volcano plot representing the HDAC6 interactome (data were extracted from Dowling et al.)[4]. The identified interaction between HDAC6 and G6PD is highlighted in blue. **f** In vitro deacetylase activity of immunopurified KDACs. Flag-tagged HDAC6, Sirt1, and Sirt2 were expressed in HEK293T cells and immunopurified using anti-Flag beads. The catalytic activity of the enzymes was verified using the commercially available fluorogenic FLOUR-DE-LYS assay. Fluorescence, measured following the deacetylation of the substrate, is plotted as a function of incubation time with the enzyme. Data show that the enzymes were purified in their active form. **g** Lysine 89 acetylated G6PD is a substrate of Sirt1 and Sirt2 in vivo. To identify deacetylases of AcK89, HEK293T cells were co-transfected with plasmids encoding AcK89 G6PD and the indicated KDACs or a control (an empty vector). Acetylation of AcK89 was detected by Western blotting using an anti-AcK antibody. Data show that Sirt1 and Sirt2 can deacetylate K89-acetylated G6PD when overexpressed in cells ($n$=2 biologically independent samples). The relatively low K89 acetylation level following co-expression with HDAC6 is explained by the negative effect of HDAC6 on amber suppression efficiency. Source data are provided as a Source Data file.

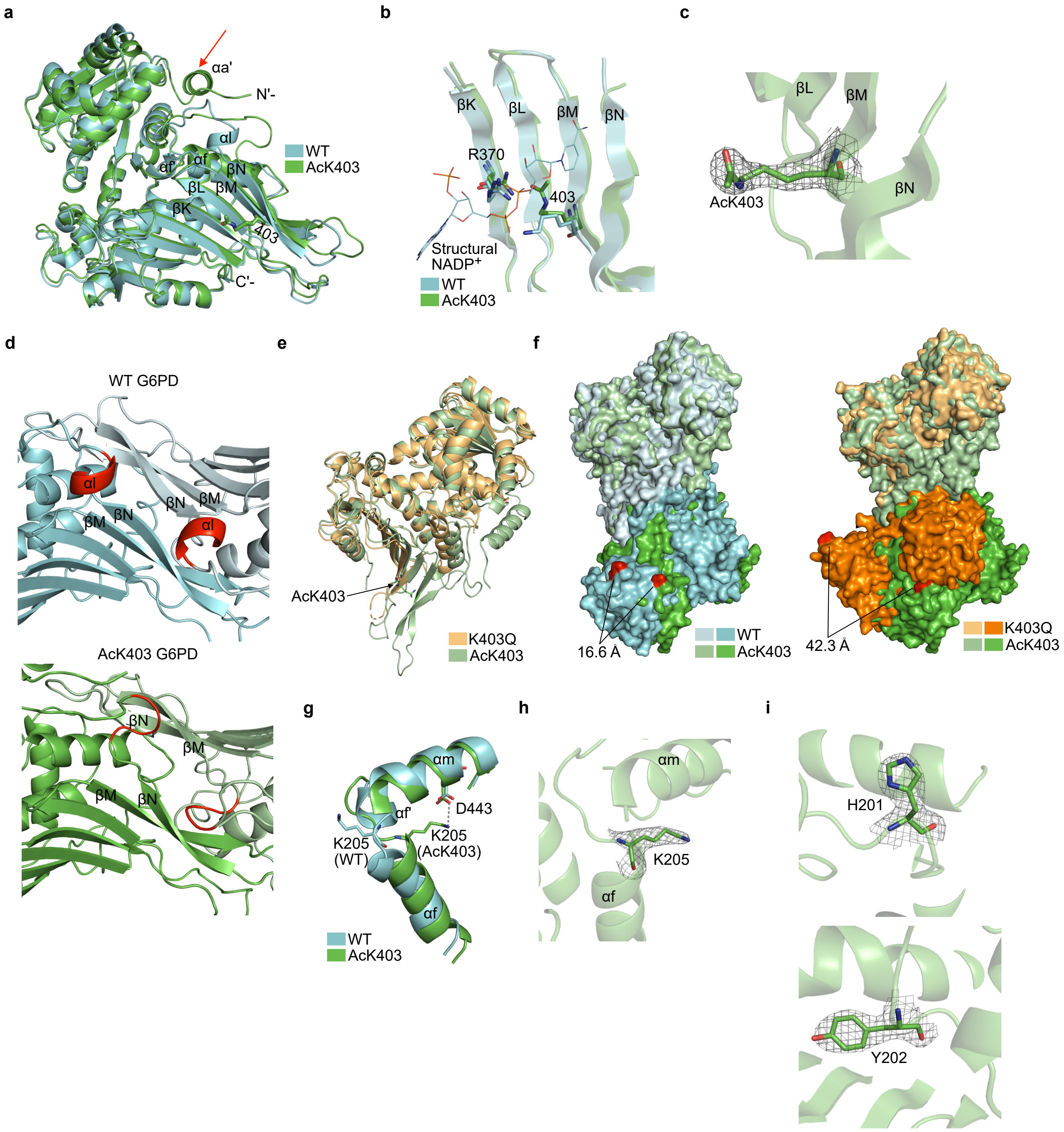


**Supplementary Fig. 7** | **a** Superposition of monomeric AcK403 G6PD (green) and WT G6PD (cyan, PDB ID: 6E08). The structure of AcK403 included an α helix at the N-terminus (helix αa', marked with a red arrow) that was not observed in other crystal structures of G6PD. **b** Superposition of AcK403 (green) and WT G6PD (cyan, PDB ID: 6E08), focusing on the structural NADP$^+$ binding site (NADP$^+$ molecule is displayed in lines). The electrostatic interaction between R370 and AcK403 is enabled by acetylation, which renders K403 neutral and probably interferes with the binding of structural NADP$^+$. **c** Electron density map around residue AcK403. The 2F$_o$-F$_c$ map was contoured at 1 σ. **d** Close-up view of part of the monomer-monomer interface around βN and αl in WT G6PD (top, PDB ID: 6E08) and AcK403 G6PD (bottom). Residues 424–427 that comprise αl in WT G6PD are highlighted in red. The short helix αl unfolds following K403 acetylation.

**Supplementary Fig. 7 Continued:** **e** Superposition of monomeric AcK403 G6PD (light green) and K403Q mutant of G6PD (light orange, PDB ID: 7SEI). Notable differences between the structures were found around the dimer interface. **f** Space-filling model of dimeric AcK403 (shades of green) compared to dimeric WT G6PD (left, shades of cyan, PDB ID: 6E08) or dimeric K403Q mutant of G6PD (right, shades of orange, PDB ID: 7SEI), with residue Glu93 highlighted in red. Structures are compared by superimposing the top monomer (light shades). The differences in the relative orientation of the lower monomers (dark shades) are highlighted by measuring the distance between residue Glu93 in AcK403 and residue Glu93 in the superimposed structure. **g** Close-up view of K205 in AcK403 G6PD (green) and WT G6PD (cyan, PDB ID: 6E08). The position of K205 in AcK403 is stabilized by electrostatic interaction with D443, which pulls together helices αf and αm. Residues comprising the short helix αf' adopt an unfolded conformation in AcK403 G6PD. **h** Electron density map around residue K205. The $2F_o$-$F_c$ map was contoured at 1 $\sigma$. **i** Electron density map around residues H201 (top) and Y202 (bottom). The $2F_o$-$F_c$ map was contoured at 1 $\sigma$.

**a**

| #1 | a⁺ | a²⁺ | b⁺ | b²⁺ | Seq. | y⁺ | y²⁺ | #2 |
|---|---|---|---|---|---|---|---|---|
| 1 | 72.08078 | 36.54403 | 100.07569 | 50.54148 | V | | | 10 |
| 2 | 200.13935 | 100.57331 | 228.13427 | 114.57077 | Q | 1129.49253 | 565.24990 | 9 |
| 3 | 297.19212 | 149.09970 | 325.18703 | 163.09715 | P | 1001.43395 | 501.22062 | 8 |
| 4 | 411.23504 | 206.12116 | 439.22996 | 220.11862 | N | 904.38119 | 452.69423 | 7 |
| 5 | 540.27764 | 270.64246 | 568.27255 | 284.63991 | E | 790.33826 | 395.67277 | 6 |
| 6 | 611.31475 | 306.16101 | 639.30967 | 320.15847 | A | 661.29567 | 331.15147 | 5 |
| 7 | 710.38317 | 355.69522 | 738.37808 | 369.69268 | V | 590.25856 | 295.63292 | 4 |
| 8 | 953.41282 | 477.21005 | 981.40774 | 491.20751 | Y-Phospho | 491.19014 | 246.09871 | 3 |
| 9 | 1054.46050 | 527.73389 | 1082.45542 | 541.73135 | T | 248.16048 | 124.58388 | 2 |
| 10 | | | | | K | 147.11280 | 74.06004 | 1 |

Y401-P, pWT G6PD:

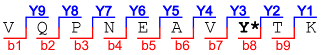


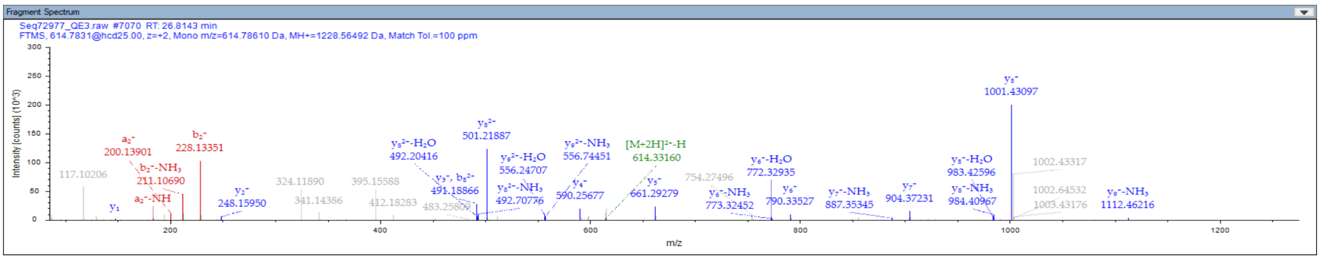


**b**

| #1 | a⁺ | a²⁺ | b⁺ | b²⁺ | Seq. | y⁺ | y²⁺ | #2 |
|---|---|---|---|---|---|---|---|---|
| 1 | 72.08078 | 36.54403 | 100.07569 | 50.54148 | V | | | 10 |
| 2 | 129.10224 | 65.05476 | 157.09715 | 79.05222 | G | 1172.46598 | 586.73663 | 9 |
| 3 | 276.17065 | 138.58897 | 304.16557 | 152.58642 | F | 1115.44452 | 558.22590 | 8 |
| 4 | 404.22923 | 202.61825 | 432.22415 | 216.61571 | Q | 968.37610 | 484.69169 | 7 |
| 5 | 647.25889 | 324.13308 | 675.25380 | 338.13054 | Y-Phospho | 840.31753 | 420.66240 | 6 |
| 6 | 776.30148 | 388.65438 | 804.29640 | 402.65184 | E | 597.28787 | 299.14757 | 5 |
| 7 | 833.32295 | 417.16511 | 861.31786 | 431.16257 | G | 468.24527 | 234.62628 | 4 |
| 8 | 934.37063 | 467.68895 | 962.36554 | 481.68641 | T | 411.22381 | 206.11554 | 3 |
| 9 | 1097.43395 | 549.22062 | 1125.42887 | 563.21807 | Y | 310.17613 | 155.59170 | 2 |
| 10 | | | | | K | 147.11280 | 74.06004 | 1 |

Y503-P, AcK403 G6PD:

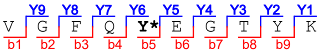


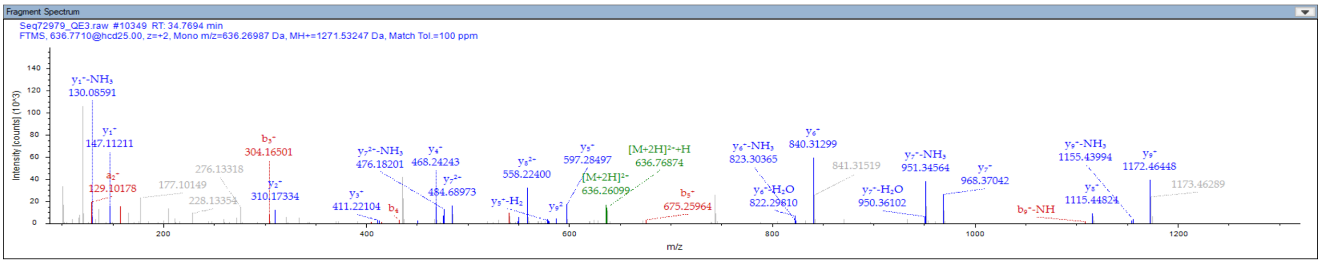


**Supplementary Fig. 8** | LC-MS/MS analyses of immunoprecipitated Flag-tagged pWT G6PD (**a**) or AcK403 G6PD (**b**) co-expressed in HEK293T cells with Fyn kinase. **Y*** marks the identified phosphorylated tyrosine residue.

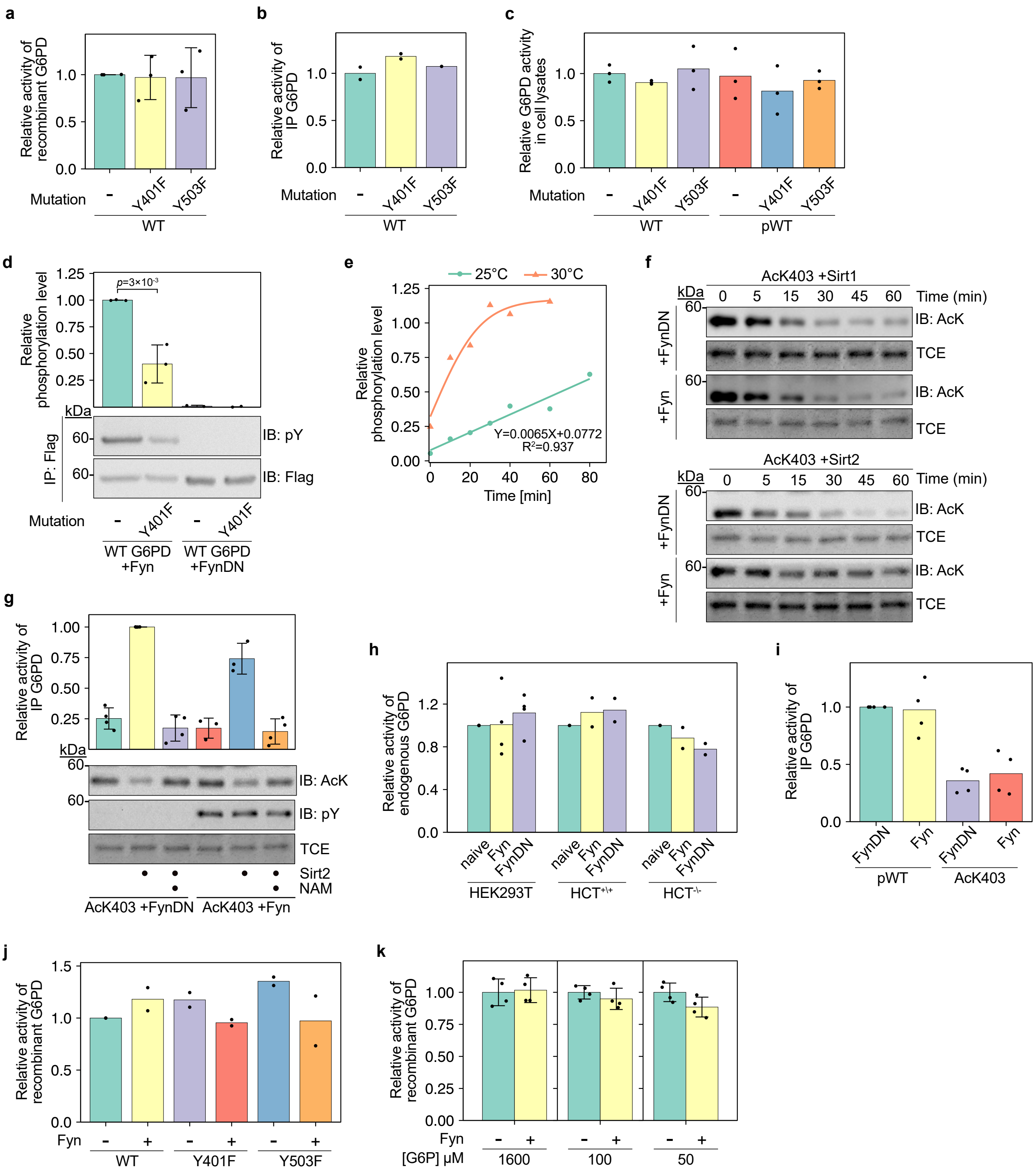


**Supplementary Fig. 9** | **a** The mutations Y401F and Y503F have no affect on $V_{\max}$ of recombinant G6PD. Bars represent the relative $V_{\max}$ of bacterially expressed and purified G6PD; $n$=3 independent experiments. **b** The mutations Y401F and Y503F have no affect on $V_{\max}$ of immunopurified exogenous G6PD. Bars represent the relative $V_{\max}$ of immunopurified Flag-tagged G6PD expressed in HEK293T cells. Enzyme activity was measured under $V_{\max}$ conditions and normalized to protein expression level; $n$=2 biologically independent samples. **c** The mutations Y401F and Y503F have no affect on $V_{\max}$ of WT and pWT G6PD in cell lysates. Bars represent the relative $V_{\max}$ of indicated G6PD variants. Protein activity was measured under $V_{\max}$ conditions in cleared cell lysates and normalized to protein expression level; $n$=3 biologically independent samples. **d** Y401 G6PD is a substrate of Fyn. HEK293T cells were co-transfected with a plasmid encoding Flag-tagged WT or Y401F G6PD and a plasmid encoding Fyn or dominant negative variant of Fyn (FynDN). The tyrosine phosphorylation level of immunoprecipitated Flag-tagged G6PD was assessed by immunoblotting. Bars represent normalized immunoblot intensities of the phospho-tyrosine (pY) antibody relative to WT G6PD co-expressed with Fyn. Data were analyzed using one-way ANOVA followed by Tukey's post hoc test and are presented as mean values ± SD; $n$=3 (+Fyn), or 2 (+FynDN) biologically independent samples.

**Supplementary Fig. 9 Continued:** **e** Time-dependent in vitro phosphorylation of recombinant WT G6PD by Fyn. Purified G6PD was incubated with 10 ng of active GST-Fyn at either 25°C or 30°C. The level of G6PD phosphorylation was monitored as a function of time by immunoblotting. In-gel TCE fluorescence was used as a loading control. Immunoblot intensities at 25°C were fitted to a linear equation (green line). The orange line (30°C) serves as a guide to the eye. **f** Fyn-dependent phosphorylation has no effect on AcK403 deacetylation by Sirt1, and Sirt2. Western blots show the acetylation level of immunoprecipitated AcK403 G6PD co-expressed in HEK293T cells with Fyn or FynDN, and incubated in vitro with Sirt1 (top) or Sirt2 (bottom) for the indicated amount of time. **g** Fyn-dependent phosphorylation does not affect the reactivation of AcK403 by Sirt2. Representative Western blots showing acetylation and phosphorylation levels of immunoprecipitated AcK403 G6PD co-expressed in HEK293T cells with Fyn or FynDN, and incubated in vitro for 1 h with or without Sirt2 and NAM. Bars represent relative $V_{\max}$ following incubation with Sirt2. Data are the mean ± SD; $n$=3 (Fyn) or 4 (FynDN) biologically independent samples. **h** Fyn-dependent phosphorylation has no effect on $V_{\max}$ of endogenous G6PD. Bars represent $V_{\max}$ of endogenous G6PD in lysates of naïve cells [HEK293T, p53 knockout HCT116 (HCT$^{-/-}$) or HCT116 (herein HCT$^{+/+}$)], or cells overexpressing Fyn or FynDN; $n$=4 (HEK293T) or 2 (HCT116) biologically independent samples. **i** Fyn-dependent phosphorylation has no effect on $V_{\max}$ of immunoprecipitated exogenous G6PD. Bars represent $V_{\max}$ of pWT or AcK403 G6PD co-expressed with Fyn or FynDN; $n$=4 biologically independent samples. **j** In vitro Fyn phosphorylation has no effect on $V_{\max}$ of recombinant G6PD. Bars represent $V_{\max}$ of purified recombinant G6PD following 40 min incubation at 25°C with (+) or without (-) Fyn; $n$=2 independent measurements. **k** In vitro Fyn phosphorylation has no effect on the activity of recombinant G6PD at low substrate concentrations. Recombinant G6PD was incubated in vitro with (+) or without (-) active Fyn for 60 min at 30°C to maximize G6PD phosphorylation by Fyn (see panel **e**). G6PD activity was measured under $V_{\max}$ conditions ([G6P]=1600 µM) as well as at substrate concentrations close to $K_{\mathrm{M}}^{\mathrm{app}}$ (Supplementary Table 2). Bars represent the relative G6PD catalytic activity measured at indicated conditions; $n$=4 independent experiments. Source data are provided as a Source Data file.

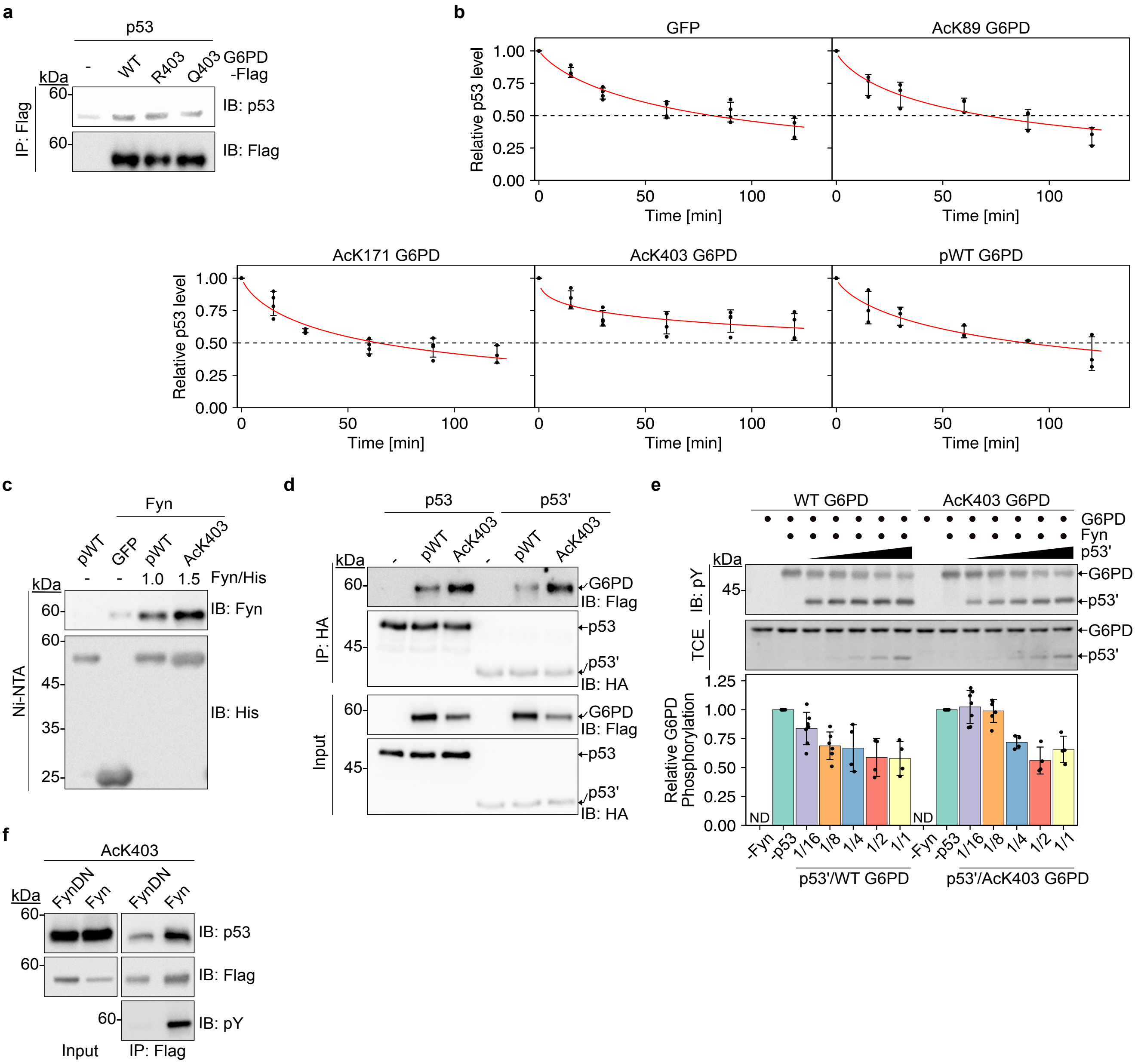


**Supplementary Fig. 10** | **a** G6PD K403Q mutation does not promote the interaction with p53. Crude extracts of HEK293T cells co-expressing full-length p53 and indicated G6PD variants were subjected to co-IP using an anti-Flag antibody. The resulting immune complexes were analyzed by Western blotting. **b** AcK403 G6PD stabilizes endogenous p53 in HCT116 cells. HCT116 cells transiently expressing the indicated G6PD variants, or GFP as a control, were treated with 50 µg/mL cycloheximide (CHX), and p53 levels as a function of time post CHX addition were evaluated by immunoblotting. Data points represent the amount of p53 relative to its level before CHX treatment; $n$=3 (pWT, AcK89, AcK171) or 4 (GFP, AcK403) biologically independent samples. Red and dashed lines serve as a guide to the eye. **c** Lysine 403 acetylation stabilized the interaction between G6PD and Fyn in bacteria. His-tagged proteins (GFP, pWT, AcK403 G6PD) and Fyn were co-expressed in CobB knockout *Escherichia coli* (*E. coli*) BL21(DE3) cells for 6 h at 37°C. Total bacterial lysates were incubated with Ni-NTA agarose beads, and co-elution of Fyn was evaluated by immunoblotting using a Fyn-specific antibody. The ratio of anti-Fyn to anti-6×His immunoblot intensities suggests a stronger interaction between AcK403 G6PD and Fyn (1.5) relative to the interaction of pWT G6PD with Fyn (1) when co-expressed in bacteria. $n$=2 biologically independent samples.

**Supplementary Fig. 10 Continued: d** AcK403 G6PD interacts with both FL p53 and p53'. HEK293T cells were co-transfected with plasmids encoding the expression of Flag-tagged G6PD and p53-HA, as indicated. Extracts were immunoprecipitated with anti-HA antibody, and immunoblotted using anti-Flag and anti-HA antibodies. **e** p53' reduces Fyn-dependent G6PD phosphorylation in vitro, but K403 acetylation mitigates the inhibitory effect at substoichiometric amount of p53'. The in vitro Fyn kinase assay was performed for 40 min at 25°C in a total volume of 20 µl, using 1.3 µM G6PD, 10 ng Fyn, and indicated amount of p53' (p53:G6PD molar ratio of 1:16–1:1). Phosphorylation products were analyzed by immunoblotting. Top: representative Western blot showing tyrosine phosphorylation level, following in vitro Fyn kinase assay in the presence of a constant amount of Fyn and G6PD, but increasing amounts of p53'. Bottom: bars represent the ratio between anti-pY immunoblot intensities and TCE quantification, relative to the ratio calculated for samples incubated without p53'; $n$=4, 6, or 7 independent experiments. In addition, the immunoblot analysis shows that Fyn phosphorylates both G6PD and p53' in vitro. **f** Fyn phosphorylation does not compromise p53 binding to AcK403 G6PD. Lysates were prepared from HEK293T cells coexpressing Fyn or FynDN with Flag-tagged AcK403 G6PD, and proteins were immunoprecipitated with anti-Flag antibody. The immunoprecipitates were subjected to Western blotting with antibodies against the Flag epitope and p53. $n$=2 biologically independent samples. Source data are provided as a Source Data file.

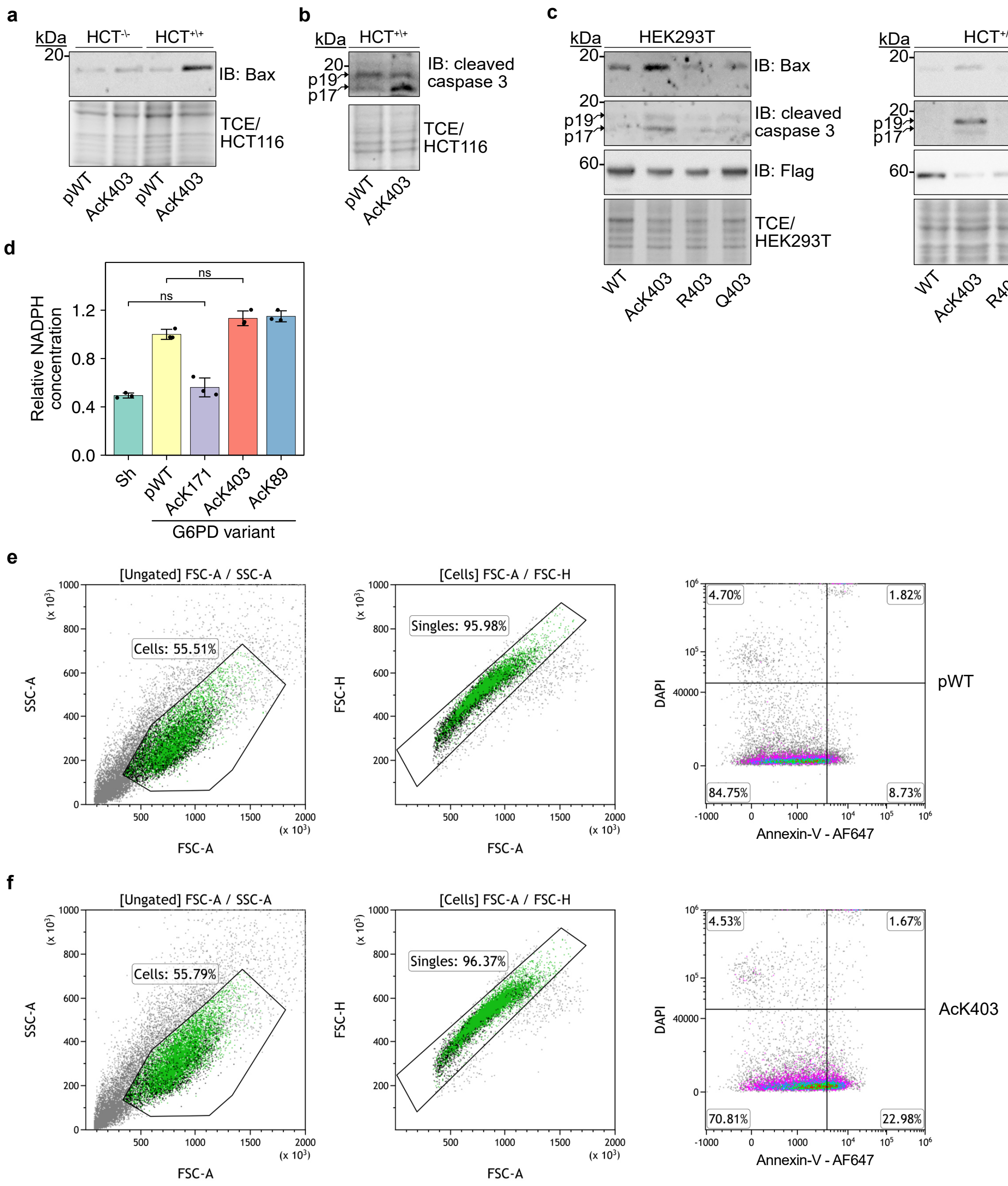

**Supplementary Fig. 11** | **a-b** Representative Western blots showing levels of endogenous Bax (a) and cleaved caspase 3 (b) in HCT116 cells. WT (HCT$^{+\backslash+}$) and p53 knockout (HCT$^{-\backslash-}$) cells were transiently transfected with plasmids encoding the expression of pWT G6PD or AcK403 G6PD. Forty-eight hours post-transfection, floating and adherent cells were collected and boiled in 1×reducing Laemmli sample buffer for 10 min. The total cell lysates were subjected to Western blot analysis using anti-Bax (a) or anti-cleaved caspase 3 (b) antibodies; total protein (visualized by TCE) served as a loading control. $n$=2 biologically independent samples. **c** Induction of pro-apoptotic signaling is dependent on K403 acetylation. Western blots show the expression levels of pro-apoptotic Bax, cleaved caspase 3, and G6PD (anti Flag) in HEK293T (left) or WT HCT116 (right) cells. Acetylation of K403 resulted in a robust increase in Bax cellular levels. Bax was also expressed at low levels following the expression of WT G6PD (mainly in HEK293T cells), but the elimination of K403 acetylation by the K403R mutation resulted in a notable decrease in Bax expression level. **d** Acetylation of G6PD does not affect the cellular concentration of NADPH. HEK293T cells cultured without KDACi and expressing the indicated G6PD variant were lysed, and NADPH concentration was measured in equal amounts of total cell lysate. Expression of inactive AcK171 G6PD had no significant effect on NADPH concentration, relative to cells expressing only sh against endogenous G6PD. The level of NADPH increased following the expression of pWT, AcK403, and AcK89 G6PD, with non-significant variability between the samples. Data were analyzed using one-way ANOVA followed by Tukey's post hoc test and are presented as mean values ± SD; $n$=3 biologically independent samples. **e-f** Representative flow cytometry analyses of cells expressing pWT (**e**) or AcK403 G6PD (**f**). Annexin V-positive and DAPI-negative cells were considered apoptotic. Source data are provided as a Source Data file.

### Supplementary Tables

**Supplementary Table 1** | Identified lysine acetylation sites in WT and site-specifically acetylated G6PD expressed in mammalian cells.

| G6PD Variant | KDACi | Position | Identified peptide | Modified PSMs | Unmodified PSMs |
|---|---|---|---|---|---|
| WT | Minus | 89 | KQSEPFFK*ATPEEK | 1 | 0 |
| | | 275 | DVMQNHLLQMLCLVAMEK*PAST NSDDVRDEK | 1 | 46 |
| | Plus | ND | ND | ND | ND |
| AcK89 | Minus | 89 | KQSEPFFK*ATPEEK | 15 | 0 |
| | Plus | 89 | KQSEPFFK*ATPEEK | 9 | 0 |
| AcK95 | Minus | 95 | ATPEEK*LKLEDFF | 30 | 1 |
| | | 97 | ATPEEKLK*LEDFFA | 1 | 29 |
| | | 293 | VLK*CISEVQANNVVLGQYVGN PDGEGEATK | 1 | 0 |
| | | 408 | MMTKK*PGMFFNPEESELDLTY GNR | 1 | 0 |
| | Plus | 95 | ATPEEK*LKLEDF | 50 | 2 |
| | | 97 | ATPEEKLK*LEDFFAR | 1 | 50 |
| | | 275 | DVMQNHLLQMLCLVAMEK*PAST NSDDVRDEK | 1 | 33 |
| AcK386 | Minus | 275 | DVMQNHLLQMLCLVAMEK*PAST NSDDVRDEK | 3 | 49 |
| | | 238 | DNIACVILTFK*EPFGTEGR | 1 | 129 |
| | | 386 | LQFHDVAGDIFHQQCK*R | 8 | 1 |
| | Plus | 97 | LK*LEDFFAR | 1 | 4 |
| | | 386 | LQFHDVAGDIFHQQCK*R | 6 | 0 |
| AcK403 | Minus | 275 | DVMQNHLLQMLCLVAMEK*PAST NSDDVRDEK | 1 | 48 |
| | | 403 | VQPNEAVYTK*MMTK | 10 | 0 |
| | Plus | 403 | VQPNEAVYTK*MMTK | 14 | 0 |
| AcK408 | Minus | 275 | DVMQNHLLQMLCLVAMEK*PAST NSDDVRDEK | 2 | 44 |
| | | 408 | K*PGMFFNPEESELDLTYGNR | 184 | 7 |
| | Plus | 275 | DVMQNHLLQMLCLVAMEK*PAST NSDDVRDEK | 2 | 50 |
| | | 408 | K*PGMFFNPEESELDLTYGNR | 207 | 0 |
| AcK432 | Minus | 275 | DVMQNHLLQMLCLVAMEK*PAST NSDDVRDEK | 1 | 253 |
| | | 407 | MMTK*KPGMFFNPEESELDLTY GNR | 2 | 2 |
| | | 408 | MMTKK*PGMFFNPEESELDLTY GNR | 2 | 2 |
| | | 432 | NVK*LPDAYER | 6 | 0 |
| | Plus | 432 | NVK*LPDAYER | 12 | 0 |

Indicated G6PD variants were expressed in mammalian cells cultured in the presence (plus) or absence (minus) of KDACi. Proteins were semi-purified by immunoprecipitation and analyzed by LC-MS/MS following in-gel trypsin digestion. Identified acetylated lysine residues are marked by *. PSM, peptide-spectrum match; ND, not detected.

**Supplementary Table 2** | Apparent steady-state kinetic parameters of WT and acetylated G6PD.

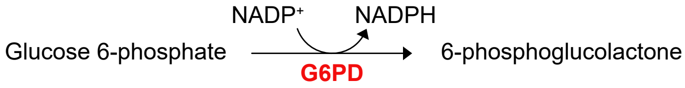


| Variant | G6P: $V_{max}$ [µM·sec$^{-1}$] | G6P: $K_M^{app}$ [µM] | G6P: $K_{cat}^{app}$ [sec$^{-1}$] | G6P: $K_{cat}^{app}/K_M^{app}$ [µM$^{-1}$·sec$^{-1}$] |
|---|---|---|---|---|
| WT | 0.053±0.002 | 62±3 | 42.4±0.6 | 0.7 |
| AcK89 | 0.102±0.004 | 124±4 | 81.8±0.6 | 0.7 |
| AcK386 | 0.031±0.002 | 45±2 | 24.5±0.3 | 0.5 |
| AcK403 | (0.002±0.001)* | (542±166)* | (1.0±0.1)* | (0.002)* |
| AcK408 | 0.0291±0.0006 | 71±3 | 23.3±0.2 | 0.3 |
| AcK432 | 0.0196±0.0004 | 80±2 | 15.7±0.1 | 0.2 |
| AcK497 | 0.026±0.001 | 56±2 | 21.0±0.2 | 0.4 |
| **Variant** | **$NADP^+$: $V_{max}$ [µM·sec$^{-1}$]** | **$NADP^+$: $K_M^{app}$ [µM]** | **$NADP^+$: $K_{cat}^{app}$ [sec$^{-1}$]** | **$NADP^+$: $K_{cat}^{app}/K_M^{app}$ [µM$^{-1}$·sec$^{-1}$]** |
| WT | 0.046±0.001 | 1.2±0.1 | 37.0±0.9 | 30.9 |
| AcK89 | 0.102±0.002 | 1.3±0.1 | 81±2 | 62.5 |
| AcK386 | 0.0281±0.0009 | 1.4±0.1 | 22.4±0.6 | 16.0 |
| AcK403 | 0.00216±0.00005 | 0.43±0.05 | 1.73±0.03 | 4.0 |
| AcK408 | 0.0294±0.0008 | 1.3±0.1 | 23.6±0.6 | 18.2 |
| AcK432 | 0.0227±0.0006 | 1.2±0.1 | 18.2±0.3 | 15.2 |
| AcK497 | 0.0249±0.0006 | 1.2±0.1 | 20.0±0.3 | 16.7 |

Values were extracted from data presented in Supplementary Fig. 4a and 4b, and summarized in Fig. 2b. Top: reported parameters were measured at increasing concentrations of G6P while keeping $NADP^+$ at saturation levels. Bottom: reported parameters were measured at increasing concentrations of $NADP^+$ while keeping G6P at saturation levels.

* Due to the low catalytic activity of AcK403 G6PD at low concentrations of G6P, the kinetic parameters could not be determined at a high confidence level and consequently were not included in our analyses.

**Supplementary Table 3** | Diffraction data collection and refinement statistics.

| Structure name | AcK89 G6PD | AcK403 G6PD |
|---|---|---|
| PDB ID | 7ZVD | 7ZVE |
| Data Collection | | |
| Beamline | DLS I03 | DLS I03 |
| Wavelength (Å) | 0.976 | 0.976 |
| Space group | F222 | P1 |
| Cell dimensions | | |
| a, b, c (Å) | 60.22, 172.22, 216.48 | 64.36, 122.61, 161.31 |
| α, β, γ(°) | 90, 90, 90 | 77.23, 80.99, 77.39 |
| Resolution range (Å) | 54.98–2.46 (2.56–2.46) | 117.38–2.28 (2.36–2.28) |
| $R_{\text{merge}}$ | 0.145 (2.399) | 0.263 (4.369) |
| $R_{\text{meas}}$ | 0.151 (2.492) | 0.286 (4.701) |
| $R_{\text{pim}}$ | 0.041 (0.684) | 0.112 (1.726) |
| $CC_{1/2}$ | 0.999 (0.588) | 0.992 (0.313) |
| Mean I/sigma(I) | 12.4 (1.1) | 5.0 (0.4) |
| Multiplicity | 13.4 (13.2) | 6.5 (7.3) |
| Completeness (%) | 99.9 (99.9) | 97 (97.3) |
| Refinement | | |
| Reflections used in refinement | 20725 (2058) | 205072 (20355) |
| Reflections used for R-free | 1037 (104) | 10314 (1039) |
| $R_{\text{work}}/R_{\text{free}}$ | 0.2077 / 0.2893 | 0.2107 / 0.2610 |
| Number of non-hydrogen atoms | 4026 | 32772 |
| macromolecules | 3932 | 32325 |
| ligands | 48 | 49 |
| solvent | 46 | 398 |
| Protein residues | 484 | 3987 |
| Average B-factor | 65.85 | 58.28 |
| macromolecules | 65.71 | 58.47 |
| ligands | 89.10 | 65.81 |
| solvent | 54.05 | 41.8 |
| R.m.s. deviations | | |
| RMSD bonds (Å) | 0.012 | 0.015 |
| RMSD angles (°) | 2.17 | 2.19 |
| Ramachandran favored (%) | 94.36 | 97.32 |
| Ramachandran allowed (%) | 4.80 | 2.28 |
| Ramachandran outliers (%) | 0.84 | 0.4 |

The values in parentheses refer to the data of the corresponding upper resolution shell. One crystal was used per data set. Data was collected at 100°K.

$R_{free}$ calculated using 5% of the reflection data chosen randomly and omitted from refinement.

RMSD values for bonds and angles are the respective root-mean-square deviations from ideal values.

**Supplementary Table 4** | Primers used in the course of this work.

| Oligo | Sequence |
|---|---|
| A. pBud-G6PD-Flag & pCDF-G6PD-His and Lysine to acetyl-lysine mutants | |
| pBud/NotI_Forward | AAGGAAAAAAGCGGCCGCATGGCAGAGCAGGTGG |
| pBud/KpnI_Reverse | CGGGGTACCGAGCTTGTGGGGGTTCACCCAC |
| Q83TAG_Forward | GTGGCTGACATCCGCAAATAGAGTGAGCCCTTCTTCAAG |
| Q83TAG_Reverse | TGAAGAAGGGCTCACTCTATTTGCGGATGTCAGCCACTG |
| K89TAG_Forward | CAGAGTGAGCCCTTCTTCTAGGCCACCCCAGAGGAGAAG |
| K89TAG_Reverse | TTCTCCTCTGGGGTGGCCTAGAAGAAGGGCTCACTCTGTTTG |
| K95TAG_Forward | AAGGCCACCCCAGAGGAGTAGCTCAAGCTGGAGGACTTC |
| K95TAG_ Reverse | GTCCTCCAGCTTGAGCTACTCCTCTGGGGTGGCCTTGAAG |
| K97TAG_Forward | ACCCCAGAGGAGAAGCTCTAGCTGGAGGACTTCTTTGCC |
| K97TAG_ Reverse | CAAAGAAGTCCTCCAGCTAGAGCTTCTCCTCTGGGGTGG |
| K171TAG_Forward | ACCGCATCATCGTGGAGTAGCCCTTCGGGAGGGACC |
| K171TAG_ Reverse | TCCCTCCCGAAGGGCTACTCCACGATGATGCGGTTC |
| K386TAG_Forward | TCTTCCACCAGCAGTGCTAGCGCAACGAGCTGGTGATC |
| K386TAG_ Reverse | TCACCAGCTCGTTGCGCTAGCACTGCTGGTGGAAGATG |
| K403TAG_Forward | CAACGAGGCCGTGTACACCTAGATGATGACCAAGAAGC |
| K403TAG_Reverse | TTCTTGGTCATCATCTAGGTGTACACGGCCTCGTTGGG |
| K408TAG_Forward | ACCAAGATGATGACCAAGTAGCCGGGCATGTTCTTCAAC |
| K408TAG_ Reverse | GAAGAACATGCCCGGCTACTTGGTCATCATCTTGGTGTAC |
| pBud/N414TAG_Forward | AAGCCGGGCATGTTCTTCTAGCCCGAGGAGTCGGAGCTG |
| pBud/N414TAG_ Reverse | CTCCGACTCCTCGGGCTAGAAGAACATGCCCGGCTTCTTG |
| K432TAG_Forward | AACAGATACAAGAACGTGTAGCTCCCTGACGCCTACGAG |
| K432TAG_Reverse | GTAGGCGTCAGGGAGCTACACGTTCTTGTATCTGTTGCC |
| K497TAG_Forward | GAGGCAGACGAGCTGATGTAGAGAGTGGGTTTCCAGTATG |
| K497TAG_Reverse | TACTGGAAACCCACTCTCTACATCAGCTCGTCTGCCTCCG |
| pCDF/NdeI_forward | GGAATTCCATATGGCAGAGCAGGTGGCCCTGAGC |
| pCDF/His_XhoI_Reverse | CCGCTCGAGTTAGTGGTGATGATGGTGATGGAGCTTGTGGGGGTTC ACCCAC |
| pCDF/N414TAG_Forward | CGGGCATGTTCTTCTAGCCCGAGGAGTC |
| pCDF/N414TAG_ Reverse | GCTCCGACTCCTCGGGCTAGAAGAACATGC |
| B. pBud-G6PD-Flag and Lysine to glutamine mutants | |
| K89Q_Forward | CAGAGTGAGCCCTTCTTCCAGGCCAC |
| K89Q_Reverse | CTTCTCCTCTGGGGTGGCCTGGAAG |
| K403Q_Forward | CAACGAGGCCGTGTACACCCAGATGATGAC |
| K403Q_Reverse | ATGCCCGGCTTCTTGGTCATCATCTGGGTGTA |
| C. pBud-G6PD-Flag and Lysine to arginine mutants | |
| K89R_Forward | GAGTGAGCCCTTCTTCCGCGCCAC |
| K89R_Reverse | CTTCTCCTCTGGGGTGGCGCGGAAGAAG |
| K95R_Forward | GCCACCCCAGAGGAGAGGCTCAAGC |
| K95R_Reverse | GAAGTCCTCCAGCTTGAGCCTCTCCTC |
| K97R_Forward | CAGAGGAGAAGCTCCGGCTGGAGGAC |
| K97R_Reverse | CAAAGAAGTCCTCCAGCCGGAGCTTCTC |
| K95/97R_Forward | CACCCCAGAGGAGAGGCTCAGGCTG |
| K95/97R_Reverse | GAAGTCCTCCAGCCTGAGCCTCTCCTC |
| K403R_Forward | CAACGAGGCCGTGTACACCCGCATGATGACCAA |
| K403R_Reverse | ATGCCCGGCTTCTTGGTCATCATGCGGGTGTACAC |
| D. pBud-G6PD-Flag & pCDF-G6PD-His and Tyrosine to Phenylalanine mutants | |
| Y401F_Forward | CCAACGAGGCCGTGTTCACCAAGAT |
| Y401F_Reverse | CTTCTTGGTCATCATCTTGGTGAACACGGCCTC |

*Continued on next page*

Supplementary Table 4 – *Continued from previous page*

| Oligo | Sequence |
|---|---|
| K403TAG Y401F_Forward | CCAACGAGGCCGTGTTCACCTAGATG |
| K403TAG Y401F_Reverse | GGTCATCATCTAGGTGAACACGGCCTCG |
| Y503F_KpnI_Reverse | CACGGTACCGAGCTTGTGGGGGTTCACCCACTTGTAGGTGCCCTCAAA CTG |
| Y503F_XhoI_Reverse | CACCTCGAGTTAGTGGTGATGATGGTGATGGAGCTTGTGGGGGTTCAC CCACTTGTAGGTGCCCTCAAACT |
| | E. pACYC-Fyn |
| NcoI_Forward | CACTACCATGGGCTGTGTGCAATGTAAGGATAAAG |
| NotI_Reverse | CACATGCGGCCGCTTACAGGTTTTCACCAG |
| | F. pBJ5-HDAC6 |
| H216A_Forward | CATTAGGCCTCCTGGACATGCCGCCCAGCACAGTCTTATGG |
| H216A_Reverse | CCATAAGACTGTGCTGGGCGGCATGTCCAGGAGGCCTAATG |
| H611A_Forward | GTCCCCCAGGACACGCCGCAGAGCAGGATGC |
| H611A_Reverse | GCATCCTGCTCTGCGGCGTGTCCTGGGGGAC |
| | G. pCDF-p53 |
| Δ66 p53_Forward | GACTAGGATCCATGCCAGAGGCTGCTCCCCCCGTG |
| Δ66 p53_Reverse | GACTACTCGAGTTAAGCGTAATCTGGAACATCGTATGGGTACATGTCT GAGTCAGGCCCTTCTGTC |
